# Anticipating complexity in the deployment of gene drive insects in agriculture

**DOI:** 10.1101/169938

**Authors:** Jennifer Baltzegar, Jessica Cavin Barnes, Johanna E. Elsensohn, Nicole Gutzmann, Michael S. Jones, Sheron King, Jayce Sudweeks

## Abstract

Insects cause substantial losses to agricultural crops each year and require intensive management approaches. Genetic pest management (GPM) has emerged as a viable, non-chemical alternative for managing insect pests. The development of engineered gene drives for agricultural use is promising, though unproven, and has the potential to impact farmers as well as broader socio-ecological systems in several ways. Drawing on lessons from the deployment of other pest control technologies, this paper considers how gene drive insects could intersect with some of the complexities that characterize agricultural systems. The development of gene drives is emerging in a landscape of pest management shaped by past and current approaches, experiences, regulations, public opinion and pest invasions. Because gene drive insects may spread well beyond their release area, stakeholder groups at different spatial scales need to be engaged in decisions about their deployment. This new paradigm both complicates and offers great promise for future pest management efforts.

## Introduction

Since the beginning of agriculture, societies have worked to protect crops from destruction by insect pests (Oerke 2006). Pest control strategies have tracked developments in both technological innovation and knowledge of pest behavior, incorporating mechanical, cultural, chemical, and biological approaches over time. Today, agricultural insect pests in conventional systems are often controlled with synthetic chemicals, combined in some cases with transgenic, insecticidal plants. Although current strategies prevent substantial losses, an estimated 30-40% of staple crops are still lost to the combined impact of insects and plant pathogens, many of which are insect-vectored (Oerke 2006). In addition to limited efficacy, pesticides are a suboptimal solution due to potential impacts on beneficial organisms, the evolution of resistance in insect populations, association with negative human health outcomes, and the economic burden placed on producers by the necessity of routine applications. Addressing some of these ecological and social concerns, genetic approaches to pest control have emerged as non-chemical alternatives for managing insect populations. Informed by the principles of evolutionary biology, genetic pest management (GPM) harnesses the mechanisms of genetic inheritance in sexually reproducing insect species to achieve either population suppression (the local elimination of a pest species or reduction in its population to below economically-relevant levels) or population replacement (the replacement of individuals in a population with non-pestilent variants) (Curtis 1985; Robinson 1998). This paper considers how a potentially revolutionary technique for GPM -- engineered gene drives -- might fit into complex socio-ecological landscapes that have been shaped by the political, economic, cultural, and environmental legacies of other pest management approaches.

### Background

Although interest in agricultural GPM has expanded with growing concerns over pesticide use, the potential for a genetic approach to insect control was first identified by Serebrovskii (1940) before the limitations of chemical control and the need for alternatives were fully realized (Robinson 1998). Since that time, scientists have worked to develop an approach to genetic control that would link desired traits, like sterility or vector incompetence, with “selfish” genetic elements that bias inheritance, driving through populations at rates higher than expected by the laws of Mendelian genetics (Burt and Trivers 2006; Gould 2008). However, attempts to appropriate naturally occurring selfish genetic elements have been limited by the difficulty of manipulating these complex systems in many organisms, including model species (Champer et al. 2016). Progress in subsequent efforts to reengineer selfish elements has likewise been slowed by their lack of flexibility and stability (Esvelt et al. 2014; Gould 2008).

Many of these limitations may be addressed by a new technique for cutting and modifying DNA based on a mechanism of adaptive immunity in bacteria, called CRISPR^1^/Cas (Horvath and Barrangou 2010). Co-opted as a gene editing tool, CRISPR/Cas can be used to alter genes in individual organisms or to create a self-replicating gene drive system that may remain active across many generations. In individual gene editing applications, the resulting molecular changes can be passed on to subsequent generations via normal inheritance, but the CRISPR/Cas machinery is not. Alternatively, in gene drive applications, genes coding for the CRISPR/Cas components necessary for gene editing are incorporated into the recipient organism's genome. Theoretically, because CRISPR/Cas acts as a selfish genetic element, copying itself onto homologous chromosomes, all offspring inherit the gene editing machinery. In contrast to individual applications, the process of gene editing is expected to endure in subsequent generations (Esvelt et al. 2014). No CRISPR/Cas gene drive has been field-tested or deployed, and applied use of the technique remains hypothetical. However, proof-of-concept has been demonstrated in laboratories for yeast (DiCarlo et al. 2015), fruit flies (Gantz and Bier 2015), and mosquitoes (Gantz et al. 2015; Hammond et al. 2016). These successes have inspired proposals to use CRISPR-based gene drives to combat insect-borne diseases, control invasive species, assist threatened species, and manage pest populations more specifically and sustainably (Esvelt et al. 2014).

The pace of “the CRISPR revolution” (Barrangou 2014) and application of this system to gene drive research has raised both excitement and alarm. Discussion about gene drives has been brought to the forefront of public and academic platforms (e.g., Barrangou 2014; Montenegro 2016), and the National Academy of Sciences, Engineering, and Medicine (NASEM) recently published a committee report on the responsible conduct of laboratory research and field releases with gene drive organisms (NASEM 2016). Nearly a decade ago, Gould (2008) anticipated the emergence of robust engineered gene drive systems and identified the need for an interdisciplinary approach to determining the suitability of gene drives for controlling particular pests. He outlined a range of genetic, evolutionary, ecological, economic, and ethical questions to be considered before gene drives or other forms of genetic control are deployed. Although the number of peer-reviewed publications on gene drives has grown precipitously over the last decade, integrated and in-depth consideration of many of the socio-cultural and political-economic dimensions of the technology remains limited. Some of these questions have received particularly little attention as they apply to agricultural applications of gene drives, partly because gene drive organisms developed to address human health and conservation problems are closer to applied use (e.g., NASEM 2016).

This paper considers some of Gould’s (2008) questions in the specific context of gene drive deployment for the control of insect pests in agriculture. We focus on the economic and socio-cultural concerns previously raised by Gould (2008) and discuss additional unknowns that might emerge from the ways in which insect biology intersects with cultures, technologies, and environments in agricultural systems. Specifically, this paper considers the identification of relevant stakeholders; the distribution of costs and benefits; potential impacts on specialized agricultural producers; trade patterns and regulations; and the dynamics of public engagement and approval. Drawing on the history, science, and in some cases, controversy, of other biologically-inspired technologies for the control of pest populations, including genetically-modified (GM) crops, releases of sterile insects, and classical biological control, we anticipate and describe several complexities that might characterize the potential deployment of gene drives in agricultural systems.

## Can relevant stakeholders be identified and consulted?

Gould (2008) indicated that a key ethical question to be considered prior to gene drive deployment is whether truly informed consent can be obtained from groups that stand to be positively or negatively impacted by the technology. This concern has been echoed in more recent calls for the involvement of public audiences in decision-making around the applied use of gene drives (e.g., Esvelt et al. 2014; Montenegro 2016; NASEM 2016). However, for agricultural gene drive systems, the question remains how to draw boundaries around relevant stakeholders in both space and time. Gene drive insects constitute an area-wide pest control strategy that is most effective when applied over large geographic areas (Alphey et al. 2013). In contrast to control options like insecticides that are adopted on a farm-by-farm basis, area-wide approaches have the potential to impact everyone within the release zone. Further, the molecular constructs in gene drive systems are designed to spread, and thus may impact additional populations over time. The unique nature of gene drive insects necessitates both big-picture and long-term thinking in the identification and engagement of stakeholders.

The stepwise eradication of the New World screwworm fly (*Cochliomyia hominivorax* Coquerel), an insect pest of cattle, from North and Central America, began in 1958 and preceded the emergence of academic and activist attention to public engagement in science. Genetic control in this case was achieved through the sterile insect technique (SIT), which involves the sterilization and release of a large number of flies that inundate the native pest population, leading females to mate with sterile males and produce no viable offspring (Davidson 2012). Stakeholder involvement in this program consisted of unidirectional public information campaigns, farmer and rancher education, and cooperative agreements between the governments of participating countries and international governing bodies (Klassen and Curtis 2005). While this program met little resistance, several controversies have emerged around projects that released GM organisms into human-inhabited environments without adequate public engagement. For example, in 2010, Oxitec, Ltd. (now a subsidiary of Intrexon Corporation) collaborated with the Mosquito Research and Control Unit of the Cayman Islands to release 3.3 million effectively-sterile male *Aedes aegypti* mosquitoes on Grand Cayman. The project engendered considerable backlash due to a perceived lack of sufficient public knowledge or regulatory oversight (Subbaraman 2011). The controversy surrounding this case led to increased public engagement in subsequent releases of GM mosquitoes by Oxitec, Ltd. in Brazil and Malaysia (Carvalho et al. 2015; Subramaniam et al. 2012).

In some cases, the inclusion of relevant stakeholders in the deployment of GM organisms has been limited by current understanding of socio-ecological systems. In 2012, when Monsanto obtained authorization from the Mexican government to cultivate GM soy in the Yucatan Peninsula, soy plants were believed to be exclusively self-pollinated (Villanueva-Gutierrez et al. 2014). The GM plants were not expected to impact insect pollinators, including honeybees, so honey-producing indigenous communities in the region were not consulted prior to planting. Studies later revealed that honeybees in the Yucatan had been visiting soy flowers and had incorporated GM soy pollen into honey marketed for export to countries in the European Union (EU). Given the EU’s strict limits on GM content in imports, indigenous producers stood to lose millions of dollars (Lakhani 2014) as their honey faced rejection in EU markets (Villanueva-Gutierrez et al. 2014). Because the Mexican constitution recognizes the autonomy of indigenous communities in decisions affecting their natural resources and grants them the right to consultation prior to the release of GM organisms that may impact them, Monsanto’s permit to cultivate GM soy in Mexico was ultimately rescinded (Vasquez 2005). Scientists still have limited understanding of the behavior and dispersal capabilities of many insect species associated with crops and no empirical insight into the behavior of gene drive insects in agricultural environments; this dearth of understanding could complicate the identification of relevant stakeholders for gene drive insects.

While the interests of indigenous groups in the honey case were protected by the Mexican constitution, other marginalized communities may not have the agency to decide which pest management solutions are acceptable or the organizational capacity to voice their opposition to approaches they deem unacceptable (Prno and Slocombe 2012). Further, groups must be able to articulate their concerns and demands in ways that encourage other actors to recognize and respond to them; an inability to do so can result in their dismissal as irrational or imbalanced (Gunningham et al. 2004). Importantly, it has been suggested that countries without centralized seed production -- typically, subsistence-based, developing countries -- could be most vulnerable to malevolent uses of gene drive technology against agricultural systems (Oye et al. 2014). The potential disparity between a community’s vulnerability to the possible negative impacts of gene drives and its voice in guiding their deployment heightens the need for mechanisms that can identify and engage underrepresented communities that stand to be affected by this technology.

## How might the costs and benefits of gene drive deployment be distributed?

Gould (2008) suggested that the utility of GPM would be determined case-by-case in the balance between the costs and benefits of genetic control for particular pests. Traditionally, economists analyze the potential adoption and benefits of new agricultural technologies with respect to their suitability in particular physical environments or crop growing zones (Renkow 1993), complexity of implementation (Foster and Rosenzweig 2010), risk preferences (Isik and Khanna 2003), and monetary costs (Duflo et al. 2011). The intensity of pest pressure, cost of alternative pest control strategies, and expected effectiveness of genetic control could also determine the potential suitability of GPM in specific contexts (Gould 2008). The scale at which these factors are considered is important, and the costs and benefits of gene drive insects may be unevenly distributed at many scales. Technological innovation in agriculture, including pest control, is heterogeneously adopted among countries, regions, and individual producers. The area-wide nature of the control potentially provided by gene drive insects further complicates patterns in the distribution of costs and benefits. Those who pay are not necessarily those who benefit, and those who may end up bearing negative consequences are not necessarily those who opted to accept the risks associated with use of the technology. Ecological, economic, and regulatory context is key in determining net effects and the profiles of winners and losers.

The burden of direct costs and the potential to capture economic benefits from the use of gene drive insects could depend on the payment structure surrounding both their development and deployment. Parasitic wasps for biological control of Asian citrus psyllid (*Diaphorina citri* Kuwayama), a vector of citrus greening disease, were freely distributed to Florida citrus producers by the Florida Department of Agricultural and Consumer Services (Alvarez, Solis, and Thomas 2015). Gene drive psyllids for population replacement have been proposed as an alternative strategy to control citrus greening disease. The Asian citrus psyllid has proven difficult to transform, and no GM psyllid line has been developed to date (CRDF 2016). However, if this project is successful, determining who will pay for the technology would be crucial. While research is currently funded by a federal grant (Turpin et al. 2012), it is unclear if direct or indirect producer contributions would be expected for any future releases. Previous area-wide campaigns such as the SIT screwworm project in the US have been funded publically, with some support from producer groups (Dyck et al. 2005)^2^. Economists have argued that because the benefits of SIT programs are broad and society-wide, central funding through general taxation may be more efficient (Mumford 2005). However, when isolated producer groups disproportionately benefit from a program and adequate organization is present (e.g., grower associations), targeted producer contributions may be practical and fair.

Gene drive insects would likely comprise an area-wide pest control strategy, as such, the extensive literature on the economic benefits of area-wide SIT programs provides a useful foundation for considering their potential impacts (e.g., Davis and Oelscher 1985, Enkerlin and Mumford 1997). While the large-scale benefits of SIT have been described, fine-scale analysis of the distribution of benefits among different types and sizes of producers is lacking. For example, average and aggregate benefits of screwworm SIT in Mexico were estimated (Davis and Oelscher 1985) and used to project similar benefits for the US and Central America (Wyss 2000, 2002), but no farm-level impact studies have examined the distribution of positive and negative impacts from this decades-long intervention. Similarly, aggregated cost-benefit projections (i.e., total benefits for all producers and consumers) were conducted for Mediterranean fruit fly (*Ceratitis capitata* Wiedemann) SIT suppression and eradication programs in Jordan and Palestine (Enkerlin and Mumford 1997). However, these projections cannot be used to examine how subgroups of individuals in different ecological regions or production categories (i.e., small or large) have benefited disproportionately, precluding a detailed understanding of how SIT can alter production landscapes. Higher-resolution studies of the impacts of area-wide control strategies would be valuable in predicting how gene drive insects might affect growers at an individual and collective level. These fine-scale economic forecasts of heterogeneity in gene drive benefits could facilitate the design of appropriate payment structures and the identification of the relevant stakeholders described in the previous section.

## How might gene drive insects impact specialized production groups?

The potential exists for specialized producers, including organic, pesticide-free, or GM-free farms, to both derive significant benefits and assume significant harm from the deployment of gene drive insects. Realized costs and benefits may depend on how GM insects are perceived by these producers and how they are classified by certification programs. Growers who hold values inconsistent with GM in general, or have consumers who do, may reject GPM outright; other growers may specifically oppose only the deployment of gene drive insects. The perspectives of individual producers may be influenced by how the technology is treated by relevant certification programs. Organic certification in many countries prohibits the use of GM technology, but the specifics vary. Australia has a zero-tolerance policy for GM material in organic fields, and revoked organic certification for one farmer’s canola fields due to contamination from a neighbor’s GM canola (Tripp 2015). In the US, organic food can contain trace amounts of GM material if the grower had no intention to introduce GM traits (National Organic Program 2016). However, demonstrating proof of intent has not been straightforward (Sudduth 2001), and exporting producers remain subject to the organic standards imposed by importing countries. Pollen flow between GM and non-GM crops is an ongoing issue, and laws regulating minimum planting distances vary by country (Beckie and Hall 2008, Greene et al. 2016). To date, these regulations have only been applied to GM crops, so it remains unclear how the presence of GM insect material -- whole bodies, body parts, or bodily fluids -- in organic products may impact access to organic certification and associated product premiums. The area-wide nature of gene drive insects may further complicate coexistence between producers who adopt the technology and those who elect or are mandated to exclude it.

On the other hand, specialized producers may experience reductions in pest pressure as a consequence of the adoption of gene drive insects by neighboring producers. This “halo effect,” in which significant benefits are derived by non-adopters, has been observed with use of GM maize and cotton expressing insecticidal endotoxins from the soil bacterium *Bacillus thuringiensis* (Bt) (Hutchison et al. 2010, Wan et al. 2012). Gene drive insects could reduce pest pressure below economic injury levels, possibly eliminating the need for chemical controls, and thereby benefiting the residue and resistance management programs for all area producers. If gene drive insects successfully manage pest populations, this may also open opportunities for the expansion of organic production into new regions or crops. In Canada, successful control of codling moth (*Cydia pomonella* Linnaeus) with an SIT program resulted in efforts to transition apple orchards to organic practices and establish an export market for organic apples from British Columbia (Bloem, Bloem, and Carpenter 2005). Deployment of gene drives for the control of spotted wing drosophila (*Drosophila suzukii* Matsumura), a significant global pest of soft-skinned fruit crops, could hold similar potential for organic markets. *Drosophila suzukii* is problematic for organic producers due to the limited number of approved and efficacious control options available to them (Van Timmeren and Issacs 2013); organic growers may thus benefit disproportionately from use of a gene drive *D*. *suzukii* through crop loss reduction. Although gene drive insects have the potential to both benefit and harm specialized producers, impacts would likely vary by crop, region, pest, and type of production system; case-by-case consideration is thus necessary prior to use. In complex and uncertain governance landscapes, specialized producers who rely on production methods that are not compatible with the use of gene drive insects deserve special consideration prior to any proposed deployment.

## How might gene drives interact with other pest management approaches?

Although often discussed in isolation, gene drive insects would be just one of many tools available for managing pest issues in agriculture. This technology would be incorporated into suites of existing techniques depending on location, crop, pest pressure, season, and year. Many crops are impacted by a multi-species pest complex, whose specific dynamics are key in the selection of appropriate control strategies. While broad-spectrum insecticides are useful as an immediate, first line of defense against insect pests, approaches that target specific pests can mitigate negative impacts on beneficial insect species. However, single-species technologies like gene drive may not be universally desired if pest complexes are a problem for one or more crop plants. The targeted nature of gene drive insects and other GPM techniques suggests that they might be most appropriate in systems where a primary pest causes the majority of crop damage, such as diamondback moth (*Plutella xylostella* Linnaeus) in brassica crops (Talekar and Shelton 1993). In crops where complexes of several pests require management, the suppression of primary pest populations with gene drive insects may increase secondary pest damage. A similar phenomenon was seen in China when the adoption of Bt cotton suppressed pink bollworm (*Pectinophora gossypiella* Saunders) populations, but led to an increase in plant bugs (Heteroptera: Miridae) (Lu et al. 2008). On larger scales, the release of gene drive insects in regions dominated by one pest species may precipitate complex shifts in pest dynamics in neighboring areas affected by multiple pest species.

Importantly, the efficacy of a gene drive insect might be enhanced when used in conjunction with other approaches. For example, it has been suggested that successful control of citrus greening would be most feasible with the concurrent deployment of GM insects for pest suppression, parasitic wasps for biological control, and GM citrus trees with disease resistance (Turpin et al. 2012; National Geographic Society 2014). In addition to interacting directly with other control strategies, gene drive insects would encounter the political and social legacies of historical pest management programs. Beginning in 1995, the US government and state of Florida attempted to eradicate the bacteria responsible for citrus canker (*Xanthomonas axonopodis* Hasse) with the compulsory removal of infected citrus trees from both commercial and residential areas (Alvarez, Solis, and Thomas 2015). Although landowners were compensated for tree removal, homeowners successfully challenged the program in court and it was halted (Alvarez, Solis, and Thomas 2015). The legal precedent set by that controversy may limit the capacity for state intervention in citrus greening control (Alvarez, Solis, and Thomas 2015). Any approach perceived to infringe on private property rights, perhaps including the area-wide deployment of gene drive insects, may face opposition by local communities.

## How might the deployment of gene drive insects impact international trade?

Agricultural production in global markets is burdened by the unintentional movement of pest species, making localized pest management important for preventing the movement of invasive species in addition to reducing crop loss. To prevent the introduction of non-native agricultural pests through trade, a range of domestic and international government and trade organizations have established sanitary and phytosanitary (SPS) regulations regarding the quality and safety of agricultural commodities. Under some SPS standards, certain insects are subject to strict quarantine measures that result in import rejection when insect presence is detected. GPM has historically helped countries regain access to the global market in the wake of imposed quarantines. For example, until 1995, fruit from Chile was subject to trade restrictions from importing countries due to Mediterranean fruit fly (*C*. *capitata*) establishment. An SIT program eradicated local populations in Chile, facilitating the reopening of the country’s fruit export market (Gonzalez and Troncoso 2007). Like SIT and other area-wide measures, gene drive insects for population suppression could significantly reduce the likelihood of insect presence in produce and thereby facilitate trade in cases where quarantine policies otherwise restrict it.

In addition to activating SPS measures regarding insect presence, gene drive insects may potentially trigger SPS measures that control the presence of GM material in food and animal feed. The risk of introducing GM insect material into agricultural shipments via the deployment of gene drive insects might vary both with the type of drive and the species targeted. This risk could be temporary in the context of population suppression drives, since the presence of gene drive insects would be transient. Drives for population replacement, which would maintain human-mediated changes in insect populations over long periods of time, present longer-term risks of contamination. Additionally, insect species that feed on the external portions of plants, such as diamondback moth, may have a lower potential of leaving GM material in the consumed portion of produce shipments, compared to species like Mediterranean fruit fly, whose larvae feed on the inside of fruit. If GM materials from gene drive insects are regulated like GM crops have been, the consequences for global trade would be significant and complex. From 2003 to 2005 alone, EU restrictions on the importation of GM products resulted in an estimated $2.87 billion reduction in agricultural export revenue for the US, Canada, and Argentina (Disdier and Fontagne 2010). The potential loss of trade markets, particularly in EU countries, has led to tepid consideration of GM crop approval in many African countries (Paarlberg 2006). GM restrictions can also shift trading patterns as product may be diverted to friendlier markets (Smyth et al. 2006). Adopters of gene drive insects would need to consider whether the potential benefits of domestic productivity and increased exportation to GM-friendly countries outweigh the costs of foregone GM-free markets.

A key uncertainty in the deployment of gene drive insects is how they will be classified by relevant governance structures at multiple scales, including international trade organizations, national laws, and certification programs. As described previously, resolving this uncertainty is also central to understanding the potential impacts of the technology on specialized producers. The ways in which accidental GM contamination has impacted trade in the past highlights the particular importance of material classification. Mexican honey containing pollen from GM soy plants was rejected in the EU because, at the time, pollen was classified as an “ingredient” of honey and there was zero tolerance for any detectable GM ingredients in food products imported into the EU. In 2014, the Court of Justice of the EU reclassified plant pollen as a “constituent” of honey (CJEU 2011). The acceptable level of GM pollen was thereby raised from zero to 0.9%, reducing the potential for future disruption in the trade of honey between Mexico and the EU (Birkman et al. 2013). Careful classification of GM insect contamination and establishment of tolerance levels will be vital in anticipating the complex ways in which gene drive insects may impact trade.

## Who grants the right to release gene drive insects?

While domestic and international regulations will fundamentally shape the ways in which gene drive insects are deployed, organizational decisions are also influenced by institutional, social, and cultural factors (DiMaggio and Powell 1983), including social prestige, the views of important constituents, and public opinion (Gunningham et al. 2004). Organizational legitimacy and survival depend upon conforming to these pressures, indicating that social factors may actually supersede the laws and regulations that govern a particular technology or company (Meyer and Rowan 1977). The history of GM soy in Mexico demonstrates that authorization granted by governments for the release of GM organisms into shared environments can be challenged by other stakeholders. Supra-legal social obligations are captured in the notion of the *social license to operate* (SLO), which is an informal, tacit agreement between a business or industry and the community in which it operates (Lacey and Lamont 2014). The SLO concept is relevant to any controversial technology or business activity (Demuijnck and Fasterling 2016) around which parties have divergent or competing interests (Lacey and Lamont 2014), including those that might characterize the deployment of gene drives to manage agricultural pests (see Kuiken (2017) for a broader discussion of SLO in regard to gene drives).

While regulatory approval is contingent upon compliance with legal mandates, SLO incorporates the expectations that various members of civil society have for the entities acting in their communities (Gunningham et al. 2004, 308). The right to release gene drive insects would require those interested in deployment to address the claims of an array of relevant stakeholders, who, as discussed previously, are difficult to identify *a priori*. Communities that could be directly impacted by a project tend to raise the greatest concerns (Demuijnck and Fasterling 2016; Hall et al. 2015), and most SLOs are thus formed at the community level, where the technological infrastructure will be located. However, SLO should be maintained at multiple scales, including the local level where the project will take place, and at industry, national and international levels (Hall et al. 2015). This would be particularly relevant for gene drive insects given their capacity for spread and the potential for complex impacts at regional and international scales. Importantly, the demands of some stakeholders may conflict with the values and economic concerns embedded in regulatory policy. Consequently, an organization’s social obligations are not necessarily synonymous with their legal obligations, and regulatory approval does not equate to social approval. This tension was recently demonstrated in GM diamondback moth research conducted at Cornell University (Harvey 2015). In 2015, Oxitec, Ltd. and Cornell University received approval from the United States Department of Agriculture (USDA) to conduct open field trials with GM diamondback moths in upstate New York (Oxitec 2016). Despite regulatory approval, and perhaps in response to disapproval by the local community, administrators at Cornell University decided against open field trials and only approved caged field trials (Cornell University 2016). Although regulatory approval was granted, social approval was incomplete.

Regulatory approval is often static, but social approval is context-specific and dynamic (Gunningham et al. 2004). GM papaya engineered for resistance to ringspot virus was adopted in Hawaii in 1998, but was not planted in other papaya-producing countries like Thailand, Venezuela, and Jamaica (Davidson 2008; Fermin and Tennant 2011; Gonsalves et al. 2007). More recently, the use of GM papaya in Hawaii has been challenged due to widespread contamination of organic papaya crops with GM papaya pollen and associated impacts on organic markets (Boyd 2008; Hewlett and Azeez 2008). Geographic and temporal variation in public acceptance and adoption of GM papaya has been shaped by differences in farmer engagement, the intensity of the pest problem, the relative importance of the crop to domestic and international markets, trade relationships, and the role of organizations that oppose the technology (Davidson 2008; Fermin and Tennant 2011). The right to release gene drive insects may similarly be influenced by these non-regulatory factors and dependent on social, cultural, and economic contexts.

## Does it matter which entities deploy gene drive insects in agriculture?

The history and discourse surrounding GM crops suggest that the nature of the organization intending to release GM organisms in the environment -- whether a for-profit company, nonprofit organization, or university -- matters in the acquisition of both regulatory and social approval. Control of agricultural biotechnology by for-profit companies and the concentration of power in an increasingly small number of transnational corporations has inspired a great deal of critique (Schurman and Munro 2010). Although many large agricultural biotechnology companies have initiatives that focus on the developing world, the profit-orientation of corporations has tended to direct substantial resources to problems experienced by large-scale, commercial producers. This sometimes results in the deployment of products that are impractical for small-scale, subsistence farmers. Proposed applications of CRISPR-based gene editing and gene drive to agriculture have also disproportionately focused on the priorities of large-scale, commercial farms, including the creation of more profitable animal livestock populations and the reversal of herbicide tolerance in weeds and insecticide resistance in crop pests (Montenegro 2016). Efforts to develop pigs with immunity to hemorrhagic virus, a problem plaguing small farms in sub-Saharan Africa and Eastern Europe, appears to be one of the few exceptions (Ainsworth 2015). Notably, this research is being conducted by the Roslin Institute, a nonprofit Scottish charity at the University of Edinburgh (Ainsworth 2015).

Given the controversies that surround corporate control of GM crops, the deployment of gene drive insects by nonprofit entities may be more likely to receive public support, since these organizations typically generate more public trust than for-profit companies (Steinberg 2006). Some nonprofit entities are explicitly motivated by humanitarian or environmental goals like sustainability or social justice, while the missions of other institutions, like public and land-grant universities, include the pursuit of education and research with relevance to public interests. However, even without a profit motive, nonprofit organizations can be susceptible to bias in the selection and design of the projects they undertake. The development and deployment of GM organisms is expensive. Any nonprofit organization involved in gene drive deployment would require capital for their projects, and might be influenced by the priorities of the philanthropic foundations and government agencies that fund their work. Additionally, as federal funding for scientific research continues to decline, university researchers increasingly depend upon funding from industry. The terms of funding contracts determine who owns the resulting research and its products and whether they will be privatized or held publically. Uneven public trust in these various organizations would combine in complex ways with the political-economic dynamics of science funding to determine which entities could successfully deploy a gene drive insect for agricultural pest management.

## Conclusion

Engineered gene drives, a potentially revolutionary technology for GPM, may impact socioecological systems in complex ways. We have emphasized some of the potential socio-cultural and political-economic dimensions of this technology, including the difficulty of identifying and engaging relevant stakeholders; unevenness in the distribution of costs and benefits for various producers; complexity in how the technology might interact with other pest management approaches and pest populations; the intricacies of international trade policies; and spatial and temporal variation in the social legitimacy of organizations that might deploy it. These issues have been problematic for many agricultural technologies in the past, and they continue to resurface in discussions over the use of biotechnology in particular. While we can draw on the history of other pest control approaches to anticipate complexities that might characterize the deployment of gene drive insects, engineered gene drives are without precedent in significant ways. Their potential to spread well beyond a release area requires thinking on larger geographic and temporal scales than are typically considered in the decision-making processes of labs, governments, corporations, or farms. This new paradigm for imagining the big-picture and long-term impacts of agricultural technologies complicates pest management decisions, but also offers great promise for revolutionizing the cultural sensitivity, economic viability, and sustainability of future pest management efforts.

## Acknowledgements

We are greatly indebted to faculty mentors who have supported the development of this article and have provided invaluable feedback, especially Fred Gould, Jason Delborne, and Zachary Brown. We also thank H.J. Burrack for helpful comments.

## Footnotes

1 CRISPR stands for “clustered regularly interspaced short palindromic repeats” (see Horvath and Barrangou 2010 for more detail).

2 For an extensive discussion on SIT public and grower program contributions and cost recovery, see For an extensive discussion on SIT public and grower program contributions and cost recovery, see Mumford (2005) and Dyck et al. (2005).

